# Strategies for motion- and respiration-robust estimation of fMRI intrinsic neural timescales

**DOI:** 10.1101/2024.04.30.590832

**Authors:** Andrew Goldberg, Isabella Rosario, Jonathan Power, Guillermo Horga, Kenneth Wengler

## Abstract

Intrinsic neural timescale (INT) is a resting-state fMRI (rs-fMRI) measure that reflects the time window of neural integration within a brain region. Despite the potential relevance of INT to cognition, brain organization, and neuropsychiatric illness, the influences of physiological artifacts on INT have not been systematically considered. Two artifacts, head motion and respiration, pose serious issues in rs-fMRI studies. Here, we described their impact on INT estimation and tested the ability of two denoising strategies for mitigating these artifacts, high-motion frame censoring and global signal regression (GSR). We used a subset of the HCP Young Adult dataset with runs annotated for breathing patterns (Lynch et al., 2020) and at least one “clean” (reference) run that had minimal head motion and no respiration artifacts; other runs from the same participants (n = 46) were labeled as “non-clean.” We found that non-clean runs exhibited brain-wide increases in INT compared to their respective clean runs and the magnitude of error in INT between non-clean and clean runs correlated with the amount of head motion. Importantly, effect sizes were comparable to INT effects reported in the clinical literature. GSR and high-motion frame censoring improved the similarity between INT maps from non-clean runs and their respective clean run. Using a pseudo-random frame-censoring approach, there was a relationship between the amount of censored frames and both the mean INT and mean error, suggesting that frame censoring itself biases INT estimation. A group-level correction procedure reduced this bias and improved similarity between non-clean runs and their respective clean run. Based on our findings, we offer recommendations for rs-fMRI INT studies, which include implementing GSR and high-motion frame censoring with Lomb-Scargle interpolation of censored data, and performing group-level correction of the bias introduced by frame censoring.

## INTRODUCTION

Intrinsic (neural) timescales (INT) reflect the time window of integration of a neuron, neuronal population, or brain region at rest. They are measured from the autocorrelation of neurophysiological signals using a range of recording methods—e.g., single-neuron recordings (Murray et al., 2014), calcium imaging (Pinto et al., 2020), electroencephalography (EEG) (Watanabe et al., 2019), resting-state functional magnetic resonance imaging (rs-fMRI) (Watanabe et al., 2019)—and are relevant to basic neural organizational structure (Murray et al., 2014), cognitive function (Cavanagh et al., 2020; Zeraati et al., 2023), and neuropsychiatric illness (Uscătescu et al., 2023; Watanabe et al., 2019; Wengler et al., 2020). INT, particularly as derived from rs-fMRI, are consequently garnering greater attention. Central to this trend is recent evidence suggesting that INT or related temporal-autocorrelation measures exhibit higher reliability and interpretability than other rs-fMRI metrics—including widely used resting-state functional connectivity measures which could be epiphenomenal to spatial and temporal autocorrelations in the fMRI signal (Shinn et al., 2023). Furthermore, INT measured using rs-fMRI have been validated against simultaneous EEG recordings (Watanabe et al., 2019) and exhibit excellent test-retest reliability (Wengler et al., 2020). While INT are likely vulnerable to artifacts similar to other rs-fMRI measures, the effectiveness of current methods for ensuring robust INT estimation is undetermined. This study aims to systematically investigate the impact of head motion and respiration artifacts on rs-fMRI INT and assess the efficacy of established denoising methods—namely high-motion frame censoring and global signal regression (GSR)— shown to minimize these artifacts in rs-fMRI functional connectivity.

Numerous studies suggest the broad relevance of INT to cognition, brain organization, and disease. First, INT is thought to reflect the strength of recurrent excitation in canonical cortical microcircuits (Chaudhuri et al., 2015), and is thus interpretable in the context of computational cognitive models and relevant to higher-order cognitive functions like working memory. Within canonical cortical microcircuits, the synaptic coupling between excitatory and inhibitory cells—and excitation-inhibition balance—determines the timescale of circuit activity (Binzegger et al., 2009; Wang, 2020; Wong and Wang, 2006). When functionally engaged, a longer timescale allows for persistent activity that outlasts its inputs (Brunel and Wang, 2001; Cavanagh et al., 2018; Cavanagh et al., 2016; Wang, 2008; Wong and Wang, 2006), a key substrate for maintaining information during delay periods in working-memory tasks (Brunel and Wang, 2001; Compte et al., 2000; Masse et al., 2019; Wang, 1999). In non-human primates, prefrontal cortex neurons with longer INT maintain working-memory representations during a delay period with greater fidelity than shorter timescale neurons (Cavanagh et al., 2018). In humans, prefrontal cortex INT lengthen during working memory, and the degree of lengthening predicts individual performance (Gao et al., 2020). In terms of organizational principles of the brain, a hierarchy of timescales has been observed in mice (Pinto et al., 2020), non-human primates (Manea et al., 2022; Murray et al., 2014; Spitmaan et al., 2020), and humans (Hasson et al., 2015; Hasson et al., 2008; Honey et al., 2012; Lerner et al., 2011; Raut et al., 2020; Stephens et al., 2013; Wengler et al., 2020). Finally, INT alterations have been found across several neuropsychiatric disorders, including psychosis (Uscătescu et al., 2023; Uscătescu et al., 2021; Wengler et al., 2020), autism (Uscătescu et al., 2023; Watanabe et al., 2019), Parkinson’s disease (Wei et al., 2023), epilepsy (Wang et al., 2021), obsessive-compulsive disorder (Xu et al., 2023), and Alzheimer’s disease (Murai et al., 2023; Zhang et al., 2023).

Despite the potential of INT as a translational tool to study brain organization, cognitive function and brain disease, the prior fMRI literature lacks systematic efforts to develop more precise and robust methods for INT estimation. This is particularly relevant for INT given theoretical work showing method-dependent biases exacerbated by data scarcity (Zeraati et al., 2022). Physiological and non-physiological artifacts are significant confounds in fMRI measures, particularly rs-fMRI, critically impacting result interpretation by reducing signal-to-noise ratio through complex mechanisms dependent upon artifact sources (Power et al., 2017b). However, the impact of these artifacts has not been considered in the context of INT estimation. Here, we focused on respiration and head motion as two such sources of artifacts. Given the sensitivity of INT estimation to data scarcity, we also examined biases induced by frame censoring as well as strategies for bias correction in group-level analyses.

Respiration significantly affects rs-fMRI BOLD signals. Primarily, respiratory events modify arterial pCO2 which directly influences cerebral blood flow, adding variance to signals everywhere in the brain (Kastrup et al., 1998). Two main respiratory patterns deviating from basic respiratory rhythms (i.e., eupnea) impact the fMRI signal: bursts and deep breaths. These two respiratory patterns exhibit distinct timescales, cardiovascular correlates, and influences on global fMRI signal (Lynch et al., 2020). Burst respiratory patterns—a consecutive tapering of respiratory depth spanning several minutes—lead to rhythmic, lagged, whole-brain spatiotemporal patterns in the fMRI signal concurrent with the burst-breath events (Lynch et al., 2020). Deep breaths—isolated breaths noticeably larger than surrounding ones—lead to prolonged brain-wide signal decreases (Lynch et al., 2020). Deep breaths can additionally induce substantial head motion and motion-induced artifacts (Power et al., 2018; Power et al., 2017b). In the context of INT, we would expect these global respiratory-related fMRI signal changes to inflate INT values, as the addition of a shared “respiratory signal” across multiple timepoints may increase fMRI signal temporal autocorrelation.

Head motion induces instantaneous, spatially patterned fMRI signal changes that are distance-dependent (Power et al., 2018; Satterthwaite et al., 2012). In particular, it creates a spurious fMRI-signal-correlation structure characterized by increased correlations between nearby brain regions and decreased correlations between distant regions (Van Dijk et al., 2012). These effects tend to be immediate and short-lasting, and scale with the degree of head displacement (Satterthwaite et al., 2013). Head motion can also bias functional connectivity results because certain populations tend to exhibit higher head motion than healthy individuals, including older individuals and those with severe psychopathology (Huijbers et al., 2017; Martin et al., 2018; Power et al., 2020; Williams et al., 2022). As such, head motion must be effectively managed to minimize artifacts that can confound individual-and group-level results. In the context of INT, in line with previous work (Uscătescu et al., 2021), we may expect head motion to inflate INT values, as these sporadic, transient, and strong fMRI signal changes could increase the time-series temporal autocorrelation via the introduction of shared “motion signal,” similar to respiration; however, in line with previous work (Wengler et al., 2020), shortening is also possible given that introduction of noise could decrease the temporal autocorrelation.

After assessing such common rs-fMRI artifacts in the context of INT, we set out to evaluate established denoising methods shown to mitigate physiological and non-physiological artifacts: high-motion frame censoring and GSR. For head motion, high-motion frame censoring strategies are broadly effective in minimizing resulting artifacts, including distance-dependent correlations (Ciric et al., 2017). We use frame censoring instead of other denoising strategies like independent component analysis and motion-based regression because they are not wholly effective in restoring volumes impacted by high motion, yielding distance-dependent covariance at these time points (Power et al., 2020; Satterthwaite et al., 2013). GSR, despite its potential limitations for rs-fMRI connectivity, has been the standout and most effective tool in reducing respiratory-pattern-induced artifacts (Power et al., 2020; Power et al., 2018; Power et al., 2017b). Overall, previous literature suggests that specific approaches taken to reduce physiological and non-physiological artifacts can be effective in rs-fMRI, but this remains to be tested in the context of INT estimation.

## METHODS

### Description of HCP sample and rs-fMRI acquisitions

The analyzed rs-fMRI data were a subset of the Human Connectome Project Young Adult (HCP-YA) dataset. Specifically, a subsample of this dataset was used (*n =* 399) for which detailed annotations on the main respiratory patterns—burst or deep breaths—have been generated (Lynch et al., 2020). Four fMRI runs (single-shot EPI with left-to-right phase encoding direction) were obtained for each participant during eyes-open-on-fixation with the following scanning parameters: repetition time (TR) = 720 ms; spatial resolution = 2 × 2 × 2 mm; time points = 1200; total acquisition time = 14.4 minutes/run. Preprocessing of the HCP data was performed using the HCP minimal preprocessing pipeline (Glasser et al., 2013) and the corresponding CIFTI data were used with cortical gray matter represented as vertices along the cortical surface and sub-cortical gray matter represented as voxels.

### Post-processing

Rs-fMRI data were further processed by nuisance regression of average white-matter signal, average cerebrospinal-fluid signal, and the six motion parameters along with their first derivatives as in previous work (Lynch et al., 2020). To evaluate the ability of GSR and high-motion frame censoring to correct for artifacts in the data, we included additional parameters in the nuisance regression step. For GSR, the average gray matter signal was additionally included. For frame censoring, individual index regressors were included for each high motion frame (framewise displacement [FD] > 0.3 mm). If frame censoring was performed, Lomb-Scargle interpolation (Lomb, 1976; Scargle, 1982) was used to estimate signal in the censored frames (to reduce errors during bandpass filtering). Lastly, bandpass filtering using an ideal rectangle window (Song et al., 2011) with a passband of 0.01–0.1 Hz was performed. The rs-fMRI data were processed in 4 ways: (1) with both GSR and frame censoring; (2) with GSR and without frame censoring; (3) without GSR and with frame censoring; and (4) without either GSR or frame censoring. In the case of high-motion frame censoring, autocorrelation functions could then be estimated in 3 ways: (1) by re-censoring (i.e., *zeroing*) the high-motion frames—as is typically done in functional connectivity (Ciric et al., 2017)—and calculating the autocorrelation function using the censored time series; (2) by re-censoring the high-motion frames and calculating the autocorrelation function from blocks of contiguous frames (i.e., *contiguous blocks*) following Raut et al. (Raut et al., 2019; Raut et al., 2020); or (3) by calculating the autocorrelation function using the Lomb-Scargle interpolated data (i.e., without re-censoring the high-motion frames; *L-S interpolation*) (Power et al., 2014). INT maps were subsequently estimated using the method described by Watanabe et al. (Watanabe et al., 2019) for the zeroing and L-S interpolation approaches and using the method described by Raut et al. (Raut et al., 2020) for the contiguous blocks approach, and then parcellated using the Glasser MMP1.0 cortical atlas (Glasser et al., 2016) and the FreeSurfer subcortical atlas (Fischl, 2012) yielding 360 cortical parcels and 19 subcortical parcels. See Figure 1 for an overview of rs-fMRI post-processing.

**Figure 1.**
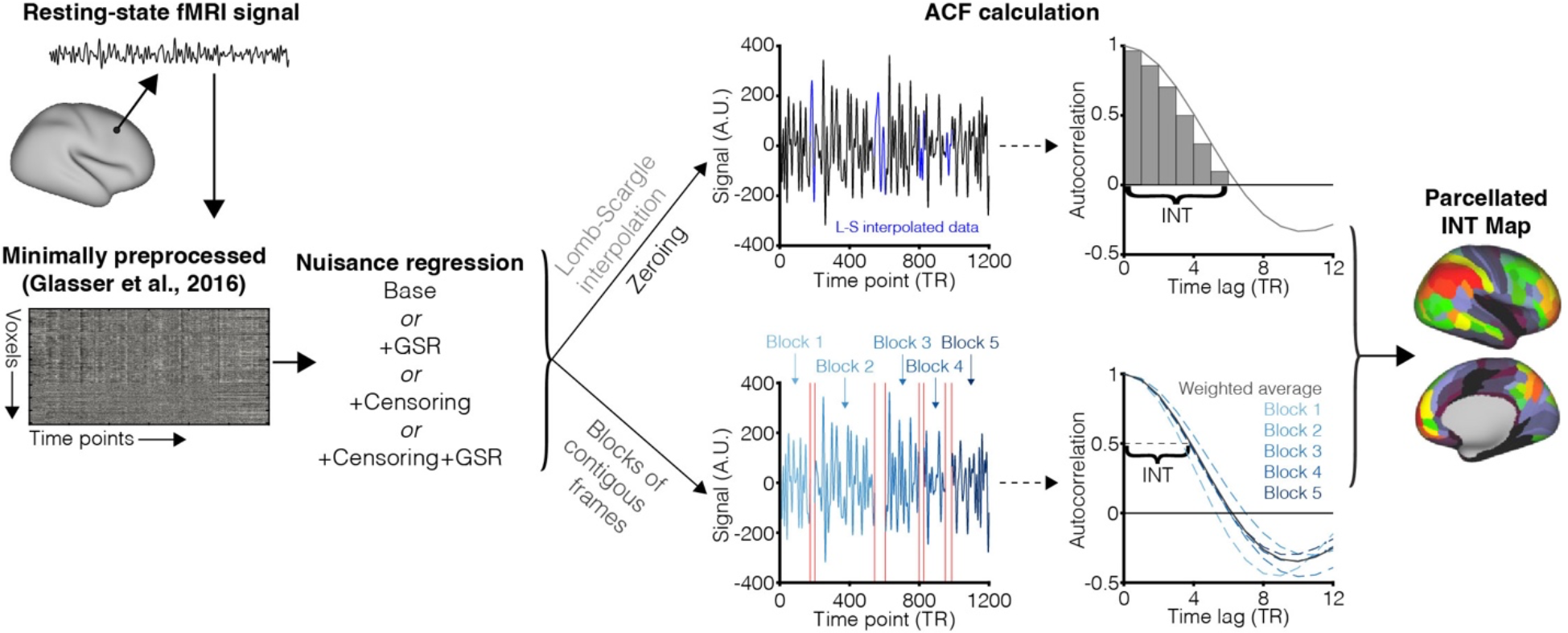
Overview of post-processing pipeline. Minimally preprocessed rs-fMRI runs from the HCP-YA dataset underwent nuisance regression for white matter (WM) and cerebrospinal fluid (CSF) signal, six motion parameters and their first derivatives, bandpass filtering in the 0.01–0.1 Hz range (i.e., “Base”) and three variations depending on the inclusion of global signal regression (GSR) and/or high-motion frame censoring. Signal autocorrelation was subsequently estimated either with Lomb-Scargle (L-S) interpolation of censored frames, no interpolation of censored frames (i.e., zeroing of censored frames), or blocks of contiguous frames (Raut et al., 2020). INT was calculated as the sum of initial non-negative period of the autocorrelation multiplied by the repetition time (Watanabe et al., 2019) for L-S interpolation and the zeroing approaches or the half-width at half-maximum (Raut et al., 2020) for the contiguous blocks approach. INT maps were then parcellated with the Glasser MMP1.0 cortical atlas (Glasser et al., 2016) and the FreeSurfer subcortical atlas (Fischl, 2012). Only cortical parcels are shown for display purposes.

### Estimation of framewise displacement (FD)

Head motion is typically calculated by first estimating changes in head position at each timepoint with six parameters (translation along the X, Y, and Z axes and rotation along the pitch, yaw, and roll axes) (Power et al., 2012). Given the high temporal resolution of the rs-fMRI HCP-YA data (i.e., TR = 0.72 s), FD was then estimated as in Power et al. (Power et al., 2019), by backwards differences over 4 timepoints, using position estimates that were filtered to suppress dominant respiratory frequencies (0.2–0.5 Hz stop band).

### Identification of “clean” and “non-clean” runs

To examine the ability of various methods to correct physiological and non-physiological artifacts, a single “clean” run deemed least likely to contain meaningful artifacts was selected and compared to other (non-clean) runs from the same individual. The rationale was that a within-subject comparison of runs with and without artifacts could help assess the ability of post-processing methods to “recover” the artifact-free clean INT data (a proxy for the ground truth) from the other artifact-ridden non-clean data, the latter of which likely comprises a substantial proportion of typical fMRI datasets and is thus more representative. The following criteria were used in conjunction to identify a run as clean: the run (1) was previously rated as not containing burst or deep breathing respiration patterns by both independent raters through visual inspection of whole-brain rs-fMRI signal plots (Lynch et al., 2020), and (2) had more than 1000 frames (out of 1200) having FD < 0.2 mm (considered a strict motion criteria). A final subset (*n* = 46) of the annotated subsample (*n* = 399) was found to have at least one clean run and used in subsequent analyses (Figure 2); for subjects with multiple clean runs (*n* = 18/46), the run with the greatest number of frames with FD < 0.2 mm was considered the clean run for that subject (the remaining clean runs were excluded from subsequent analyses detailed below). Any other runs for these 46 subjects were considered “non-clean” runs. Non-clean runs (n = 113) were divided into subtypes according to the presence of respiration patterns [deep breaths (n = 36), bursts (n = 30), or deep breaths *and* bursts (n = 20)] and degree of head motion [low motion (15% of “non-clean” runs with the lowest mean FD; n = 17); high motion (15% of “non-clean” runs with the highest mean FD; n = 17)].

**Figure 2.**
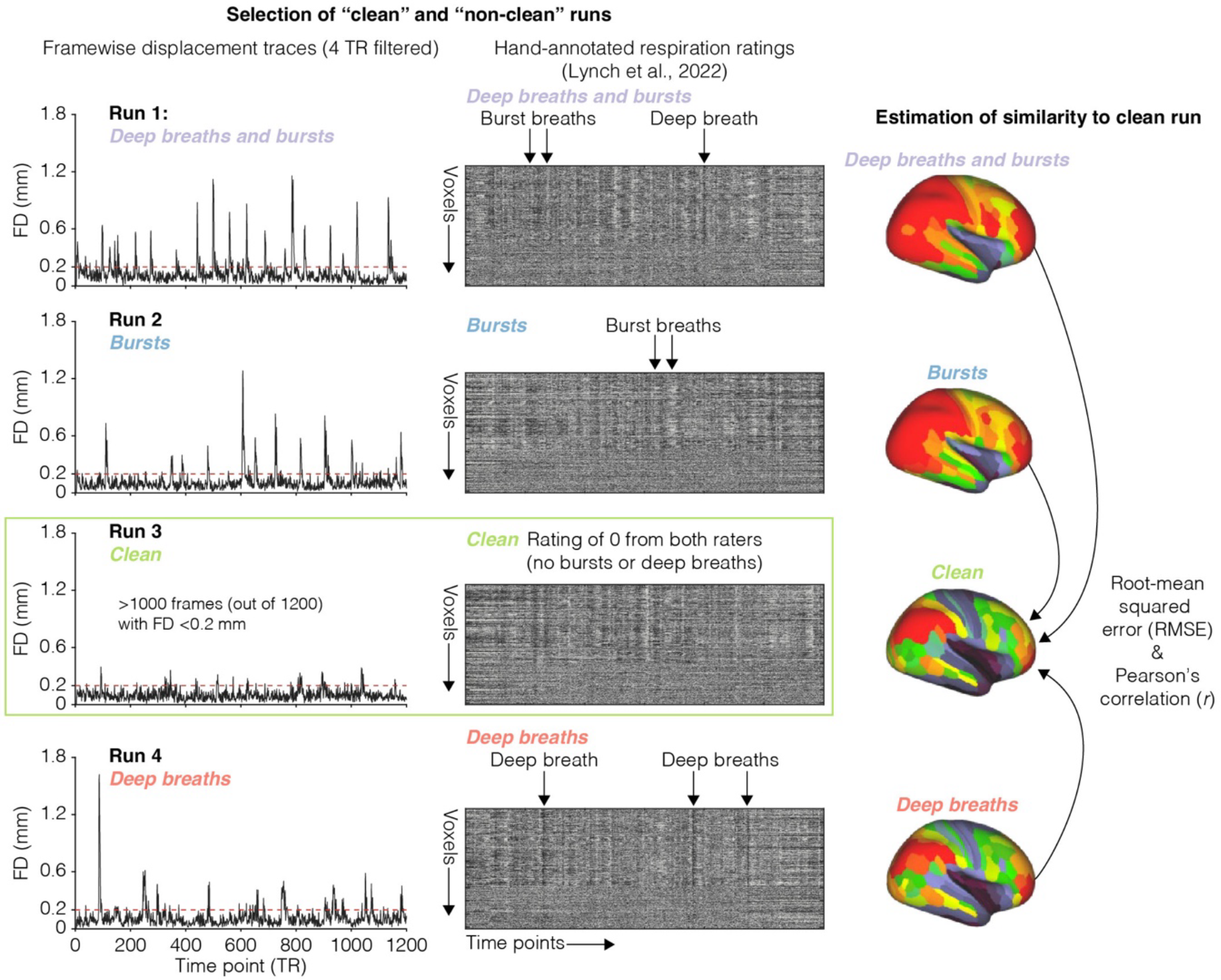
Overview of within-subject comparison approach. Motion traces and rs-fMRI gray plots (for gray matter voxels only) are shown for all four runs from one subject, each with distinct designations of clean and non-clean runs. A clean run (green box) is identified based on the amount of low-motion frames and respiration annotations from (Lynch et al., 2020). Locations of burst and deep breath patterns [from (Lynch et al., 2020)] are noted in the rs-fMRI gray plot using arrows. In analyses, the similarity between non-clean and clean runs using a within- subject comparison approach is determined by calculating the root-mean squared error and Pearson’s correlation. Exemplary parcellated whole-brain INT maps are shown. Only cortical parcels are shown for display purposes.

### Evaluation of similarity between clean runs and non-clean runs

The similarity between INT maps estimated from non-clean runs and their corresponding clean run was calculated over parcels (*p*) in two ways (Figure 2): (1) as the root-mean squared error (RMSE):

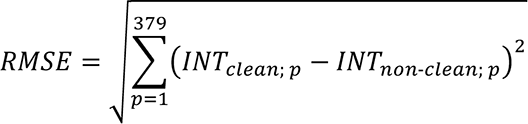

And (2) as the Pearson correlation *(r)*:

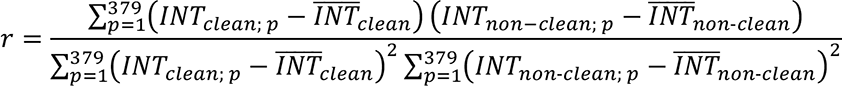

### Pseudo-random frame censoring to assess bias

To investigate the effect of frame censoring—and generally the amount of available data—on INT estimation, and our ability to correct for these effects, one analysis focused on the effects of varying levels of frame censoring, where the censored frames were pseudo-random and not based on the subject’s head motion. First, FD traces from 5 rs-fMRI runs from HCP-YA subjects not included in our subset of 46 subjects were selected that had exactly 120, 240, 360, 480, and 600 high-motion frames (FD > 0.3 mm) corresponding to 10%, 20%, 30%, 40%, and 50% frame censoring, respectively. Second, the 46 clean runs were processed as described previously, using these FD traces from these other HCP-YA subjects’ runs to select frames for frame censoring in the clean runs. We consider this process to be pseudo-random because the frame censoring is not based on true high-motion frames in the data under study but it reproduces the temporal structure of true high-motion frames observed in other data. This served to test the impact of different levels of frame censoring such as that expected when censoring high-motion frames but isolating the effect of frame censoring in the absence of concomitant head motion.

Last, the similarity between the INT maps estimated from the pseudo-randomly frame censored data and the INT maps from the non-frame-censored data was estimated as described previously.

### Group-level correction of frame censoring and motion effects

A group-level correction procedure was implemented to assess our ability to correct for bias induced by frame censoring and any residual effects of head motion not accounted for by nuisance regression. Here, we used multiple regression across subjects for a given parcel including main effects for the percentage of censored frames (*PCF*), the square of the percentage of censored frames (*PCF^2^*), and mean FD in the non-high-motion frames (*FD*; i.e., residual motion):

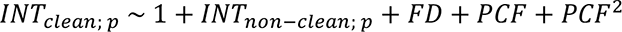

Similarity between the predicted INT values from the above model and models excluding *FD*, *PCF*, and/or *PCF^2^* and the INT maps from clean runs were estimated as described previously.

### Statistical analyses

#### Impact of respiration and head motion on INT estimation

The following linear mixed-effects (LME) models were used to assess the impact of physiological and non-physiological artifacts on INT estimation (in Wilkinson notation (Wilkinson and Rogers, 1973)):

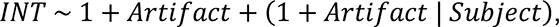

where *INT* is the average INT for each run; A*rtifact* is a dummy variable for clean and non-clean (or a type of non-clean, e.g., deep breaths) runs with clean runs as the reference; and *Subject* is an index for each subject. Random intercepts and random effects for A*rtifact* were included to account for the fact that each subject could contribute multiple non-clean runs. Bonferroni correction was used to control for multiple comparisons (n=6 for six different non-clean types).

The following LME models were used to assess the impact of head motion on INT estimation:

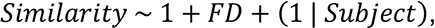

where *Similarity* is the similarity (RMSE or Pearson’s *r*) between each non-clean run and its respective clean run; and *FD* is the mean FD in the non-clean run.

#### Impact of pseudo-random frame censoring on INT estimation

The following LME models were used to assess the impact of pseudo-random frame censoring on INT estimation:

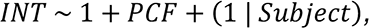

where *INT* is the average INT value for each pseudo-randomly frame censored run for a given ACF estimation method (zeroing, blocks of contiguous frames, or Lomb-Scargle interpolation); and *PCF* is the percentage of pseudo-randomly censored frames (10%, 20%, 30%, 40%, or 50%). Bonferroni correction was used to control for multiple comparisons (n=3 for three different ACF estimation methods).

The following LME models were used to assess the bias introduced by pseudo-random frame censoring to INT estimation:

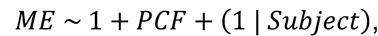

where *ME* is the mean error between each pseudo-randomly frame censored run and its respective clean (i.e., non-censored) run for a given ACF estimation method. Paired t-tests were used to compare mean error between ACF estimation methods. Bonferroni correction was used to control for multiple comparisons (n=6 for three different ACF estimation methods [LME models] and three comparisons between them [paired t-tests]).

#### Performance of GSR and high-motion frame censoring

The following LME models were used to assess the ability of GSR and high-motion frame censoring to reduce the impact of physiological and non-physiological artifacts on INT estimation:

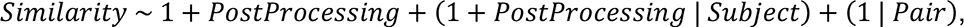

where *Similarity* is the similarity (RMSE or Pearson’s *r*) between each non-clean run and its respective clean run; *PostProcessing* is a dummy variable coding two of the four post-processing approaches (base method, including GSR, including high-motion frame censoring, and including both GSR and high-motion frame censoring) with the reduced method as the reference (e.g., including GSR for the comparison between including GSR and including both GSR and high-motion frame censoring); *Pair* is an index variable indicating each non-clean run pair. Random intercepts and random effects for *PostProcessing* were included to account for the fact that each subject could contribute multiple non-clean runs while additional random intercepts for each non-clean run pair provide equivalence to a paired t-test. Bonferroni correction was used to control for multiple comparisons (n=3 for three pair-wise comparisons with the base post-processing approach and n=2 for the comparison between including high motion frame censoring and including both GSR and high-motion frame censoring).

#### Group-level correction of bias introduced by high-motion frame censoring

Paired t-tests were used to compare the ability of the different group-level regression models (intercept only; intercept and linear frame censoring term; intercept, linear frame censoring term, and quadratic frame censoring term) to improve the similarity (RMSE and Pearson’s *r*) between each group-level-corrected pseudo-randomly frame censored run and its respective clean (i.e., non-censored) run. Bonferroni correction was used to control for multiple comparisons (n=3 for three pair-wise comparisons between the three group-level correction models).

One-sample t-tests were used to assess the residual bias (mean error) in the group-level-corrected pseudo-randomly frame censored runs for the different group-level regression models; two-sample t-tests were used to compare the residual bias between the different group-level regression models. Bonferroni correction was used to control for multiple comparisons (n=6 for one-sample tests for each of the three group-level correction models and three pair- wise comparisons between the three group-level correction models).

Paired t-tests were used to compare the ability of the different group-level regression models (intercept only; intercept and mean FD; intercept, linear frame censoring term, and quadratic frame censoring term; intercept, mean FD, linear frame censoring term, and quadratic frame censoring term) to improve the similarity (RMSE and Pearson’s *r*) between each group-level-corrected non-clean run and its respective clean run. Bonferroni correction was used to control for multiple comparisons (n=3 for three pair-wise comparisons with the intercept-only model and n=2 for the comparison between the intercept, linear frame censoring term, and quadratic frame censoring term model and the intercept, mean FD, linear frame censoring term, and quadratic frame censoring term model).

## RESULTS

### Impact of respiration and head motion on INT estimation

As expected, a within-subject comparison of clean and non-clean runs demonstrated that motion and breathing artifacts (in non-clean runs) systematically impact INT estimation (with respect to the reference clean runs for a given subject). To separately evaluate the effects of respiration and head motion, we compared clean runs to different types of non-clean runs: all non-clean runs, non-clean runs with burst breaths only, non-clean runs with deep breaths only, non-clean runs with both burst and deep breaths, and non-clean runs with either low (lowest 15%) or high (highest 15%) motion. To visualize artifacts in INT estimation spatially, for each subject with a non-clean run of a given type we calculated the difference from that subject’s clean run, and the resulting ΔINT maps for each subject were then averaged across subjects (Figure 3A). Notably, non-clean runs with burst breaths only appear to have INT alterations that occur in a particular functional pattern existing in resting-state network distribution that could reflect slow, sustained activity specific to that form of respiration (Lynch et al., 2020; Power et al., 2020). Although this is outside the scope of the current work, future studies with larger samples for each non-clean type should investigate more clearly the relationships between INT and associated neurocognitive states.

**Figure 3.**
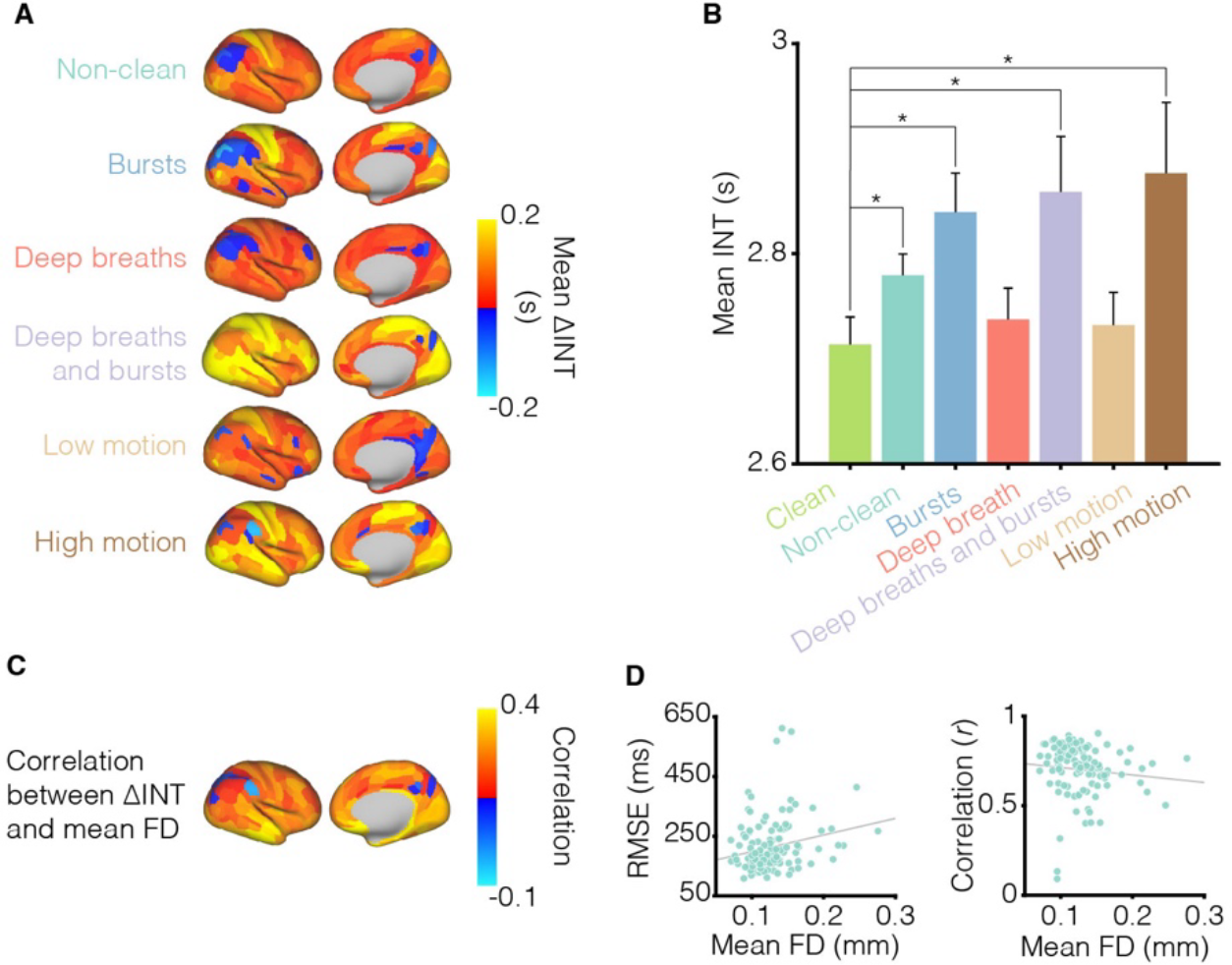
Physiological and non-physiological artifacts result in inflated INT estimates. (A) Group-averaged ΔINT (non-clean minus clean) maps for all non-clean runs and non-clean runs separated into distinct physiological and non-physiological artifact types. Non-clean runs generally exhibit elevated INT when compared to their respective within-subject clean run, although some spatially localized reductions are evident. Low motion (but still non-clean) runs exhibit near-zero changes. Only cortical parcels are shown for display purposes. (B) Group-averaged INT estimates reflect INT increases in the non-clean, bursts, deep and bursts, and high motion artifact types. (C) Increasing mean FD generally correlates with increased ΔINT across the brain. Only cortical parcels are shown for display purposes. (D) Non-clean runs with more motion had reduced similarity to their respective clean run.

The combined non-clean runs (ΔINT = 58.3 ± 125.2 ms [mean ± standard deviation]; *t*157 = 3.68, *P*Bonferroni = 0.0019), burst (ΔINT = 64.9 ± 154.7 ms; *t*74 = 3.93, *P*Bonferroni = 0.0011), both burst and deep breaths (ΔINT = 133.1 ± 116.3 ms; *t*64 = 5.10, *P*Bonferroni < 0.0001), and high motion (ΔINT = 112.1 ± 119.0 ms; *t*61 = 3.37, *P*Bonferroni = 0.0078) runs showed general increases in average INT across the brain compared to the clean runs (Figure 3B). In contrast, the deep breath (ΔINT = 44.2 ± 106.9 ms; *t*80 = 1.87, *P*Bonferroni = 0.3894) and low motion (ΔINT = 56.0 ± 130.5 ms; *t*61 = 1.51, *P*Bonferroni = 0.8188) runs did not show significant differences in average INT compared to the clean runs (Figure 3B). To further explore the effect of head motion on INT, we assessed the correlation between ΔINT (difference from clean-run INT) across the brain for non-clean runs as a function of their respective mean FD (Figure 3C). Non-clean runs with more motion were more dissimilar to the clean run from the same subject (relationship between mean FD and RMSE: *t*111 = 2.70, *P* = 0.0081; relationship between mean FD and Pearson’s *r*: *t*111 = -2.50, *P* = 0.0140; Figure 3D). Lastly, to better characterize the distinct effect of head motion aside from respiration, we identified 27 non-clean runs that were not impacted by respiration (i.e., respiration ratings of zero from both raters (Lynch et al., 2020), but < 1,000 frames with FD < 0.2 mm; “motion only”). In the motion-only runs, numerical effects were observed in the same direction that did not reach statistical significance (relationship between mean FD and RMSE: *t*25 = 1.41, *P* = 0.1699; relationship between mean FD and Pearson’s *r*: *t*25 = -1.13, *P* = 0.2708), likely due to the small sample size. Taken together, these results illustrate that respiration and head motion produce systematic and substantial biases (∼44–133 ms) in INT estimates that are comparable in magnitude to diagnostic differences reported for neuropsychiatric disorders (e.g., ∼40–90 ms difference in Alzheimer’s disease (Zhang et al., 2023), ∼40 ms difference in schizophrenia (Wengler et al., 2020)).

### Impact of high-motion frame censoring on INT estimation

Given the established utility of high-motion frame censoring to combat the effects of physiological and non-physiological artifacts, and the potential biases of data paucity on INT estimation, we sought to determine if frame censoring itself impacted INT estimation (even in the absence of large head motion) as well as the robustness of different estimation methods to such impact. To this end, we deployed pseudo-random censoring of frames (irrespective of their degree of motion) and subsequently estimated the autocorrelation in three ways: zeroing high-motion frames (Ciric et al., 2017), only using contiguous blocks of non-high-motion frames (Raut et al., 2020), and using Lomb-Scargle (L-S) interpolation of the censored high-motion frames (Power et al., 2014). Across all three methods, increasing amounts of pseudo-randomly censored frames led to decreased INT throughout the brain (Figure 4A). Increasing amounts of pseudo-randomly censored frames resulted in decreased mean INT values (zeroing: *t*228 = - 55.08; contiguous blocks: *t*228 = -24.18; L-S interpolation: *t*228 = -17.86; all *P*Bonferroni < 0.0001, Figure 4B) and greater negative mean error relative to clean runs (zeroing: *t*228 = -55.08; contiguous blocks: *t*228 = -24.18; L-S interpolation: *t*228 = -17.86; all *P*Bonferroni < 0.0001, Figure 4B). Among the methods, the L-S interpolation method was most robust, as it generally exhibited less negative mean error compared to both the zeroing method (*t*229 = 25.20, *P*Bonferroni < 0.0001; Figure 4B) and the contiguous blocks method (*t*229 = 33.74, *P*Bonferroni < 0.0001; Figure 4B). The contiguous blocks method also resulted in less mean error than the zeroing method (*t*229 = 16.39, *P*Bonferroni < 0.0001; Figure 4B). These results suggest that frame censoring itself biases INT estimation towards shorter values, and that Lomb-Scargle interpolation of the signal at high-motion frames appears most robust to this bias.

**Figure 4.**
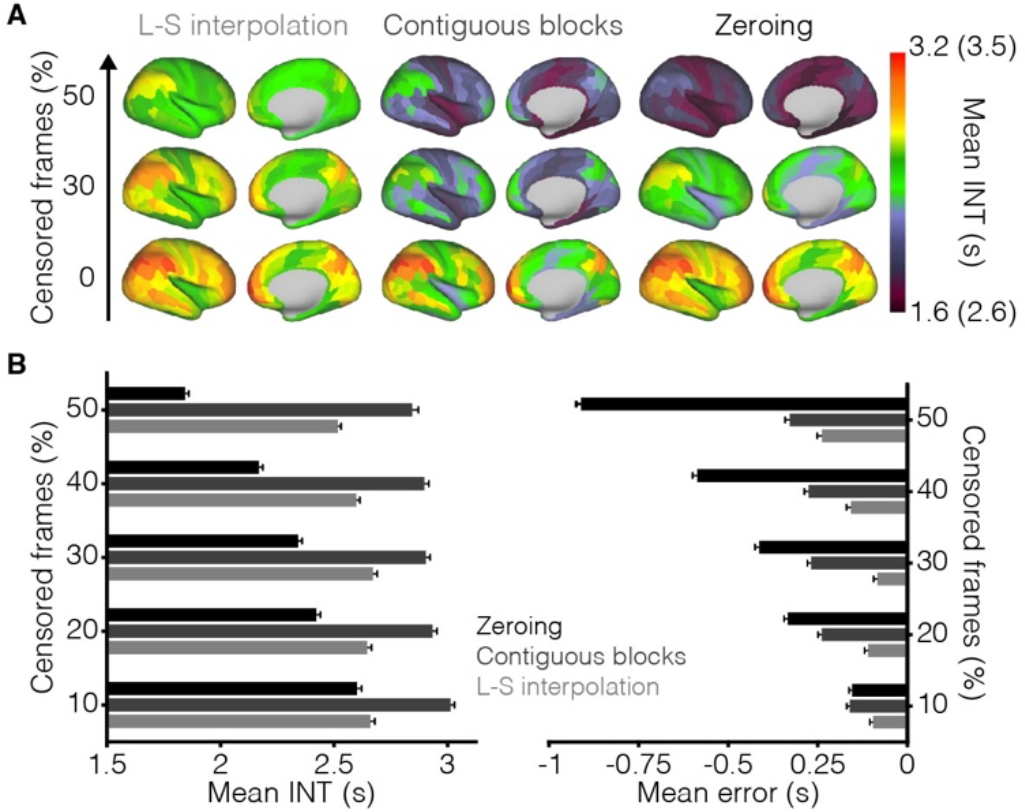
Frame censoring biases INT estimates towards shorter values. (A) Group-averaged INT maps for varying levels of pseudo-random frame censoring. Increasing pseudo- random frame censoring corresponds to brain-wide decreases in INT across the three ACF estimation methods. L-S interpolation appears most robust to this effect while zeroing censored frames was highly susceptible as expected. Only cortical parcels are shown for display purposes. A different color scale (limits in parentheses) was used for Contiguous Blocks given the slightly different INT estimation approach (see Fig. 1). (B) All three ACF estimation methods had significant linear relationships between the percentage of pseudo-randomly censored frames and whole-brain mean INT (left) and whole-brain mean error (right). The strongest relationships were for the zeroing method, followed by the contiguous blocks method, and the L-S interpolation method having the weakest (but still significant) relationships.

### Performance of GSR and high-motion frame censoring

Next, we evaluated the performance of GSR and high-motion frame censoring in the INT maps calculated using Lomb-Scargle interpolation of censored frames. For context, given the above results using pseudo-random frame censoring of up to 50% of frames, the average (± standard deviation) percent of censored frames in the non-clean runs was 4.36% ± 6.75% (range: 0%– 38.13%). Prior to either GSR or frame censoring, there is evident heterogeneity in the spatial pattern of INT between mean clean-run INT maps and those for each of the non-clean run subtypes (Figure 5A). However, the addition of GSR and frame censoring, especially together, reduces this error, as is apparent in the increased similarity between non-clean runs and the clean runs (for which the different denoising strategies unsurprisingly have little effect) (Figure 5A). Compared to the base preprocessing, frame censoring improved the similarity between non-clean runs and their respective clean run (RMSE: *t*224 = -4.85, *P*Bonferroni < 0.0001; Pearson’s *r*: *t*224 = 3.73, *P*Bonferroni = 0.0007; Figure 5B) as did GSR and frame censoring together (RMSE: *t*224 = -7.21, *P*Bonferroni < 0.0001; Pearson’s *r*: *t*224 = 4.50, *P*Bonferroni < 0.0001; Figure 5B). Although GSR on its own did not improve similarity compared to the base preprocessing (RMSE: *t*224 = - 0.44, *P*Bonferroni = 1; Pearson’s *r*: *t*224 = -0.65, *P*Bonferroni = 1; Figure 5B), it did improve similarity when combined with frame censoring when compared to frame censoring alone (RMSE: *t*224 = - 4.12, *P* = 0.0001; Pearson’s *r*: *t*224 = 2.59, *P*Bonferroni = 0.0203; Figure 5B). Together, these results support the use of both high-motion frame censoring with Lomb-Scargle interpolation of censored frames and GSR in post-processing pipelines for INT estimation from rs-fMRI data.

**Figure 5.**
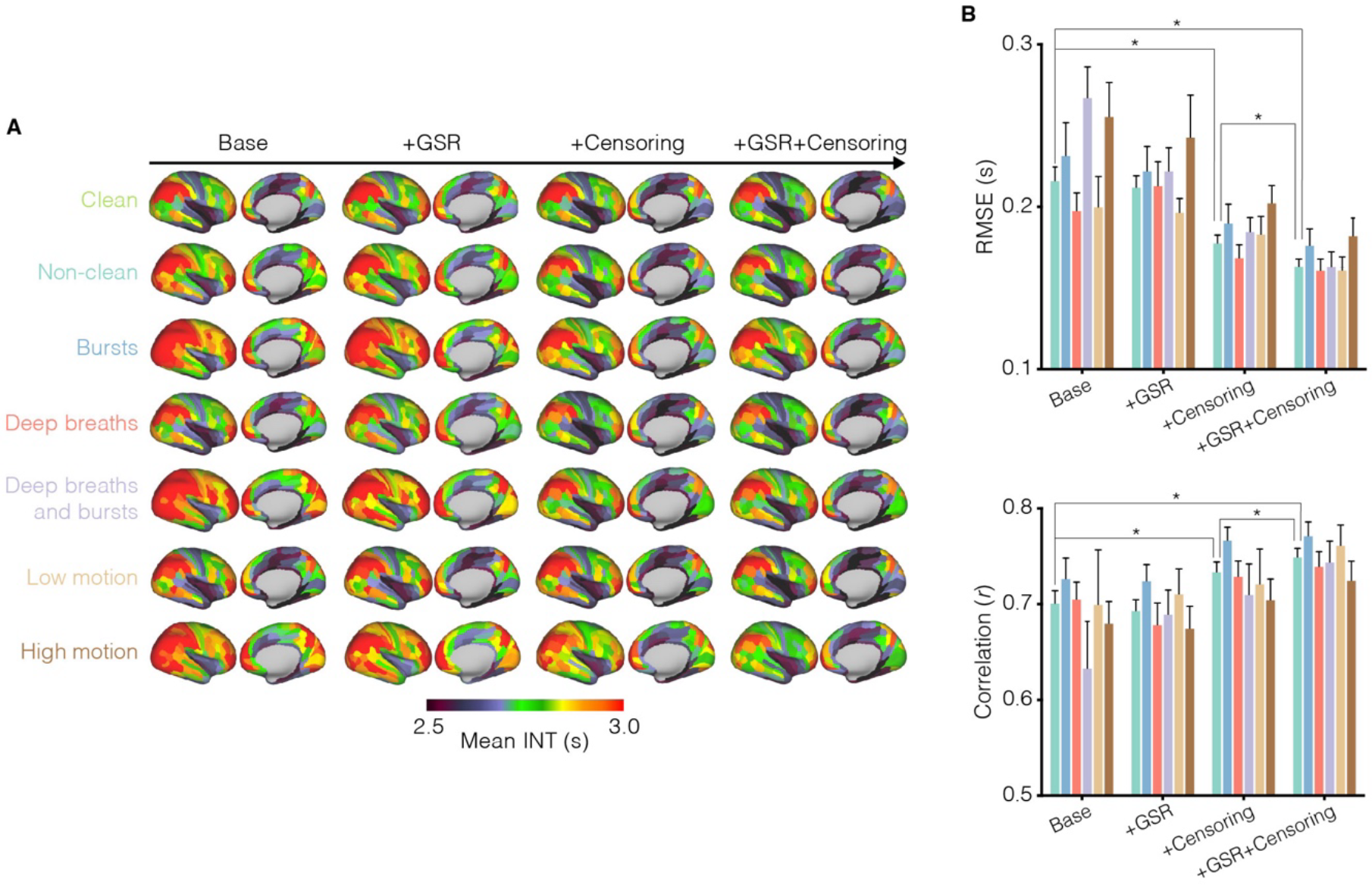
High-motion frame censoring and GSR help resolve the effects of physiological and non-physiological artifacts. (A) Whole-brain group-averaged INT maps for the base preprocessing method (leftmost column), the base method plus GSR (left-center column), the base method plus frame censoring with L-S interpolation (right-center column), and the base method plus GSR and frame censoring with L-S interpolation (rightmost column). Non-clean INT maps with the base preprocessing method exhibit significant heterogeneity, with INT elevations clearest in the non-clean, bursts, deep and bursts, and high motion artifact types (leftmost column). The addition of frame censoring and GSR improves similarity to clean maps (rightmost column). Only cortical parcels are shown for display purposes. (B) Frame censoring alone and GSR plus frame censoring significantly improves the similarity between non-clean and clean runs compared to the base preprocessing method. Furthermore, the addition of GSR to frame censoring significantly improves the similarity between non-clean and clean runs compared to frame censoring alone.

### Group-level correction of bias introduced by high-motion frame censoring

The previous results show beneficial effects of frame censoring on the accuracy of INT estimation (i.e., for yielding INT values from non-clean runs that better match those in clean runs from the same subjects; Figure 5), while also showing that frame censoring itself introduces a systematic bias towards INT underestimation (Figure 4). We next set out to test a regression-based bias-correction strategy to obtain the beneficial effects of frame censoring while minimizing the underestimation bias associated with it. We specifically tested whether a group-level correction model would improve the similarity of non-clean runs to their respective clean run. First, we attempted this analysis in the pseudo-randomly frame censored data. The inclusion of a first- and second-order term for the percentage of censored frames significantly reduced RMSE between the corrected pseudo-randomly frame censored INT maps and the clean INT maps (first-order term: *t*45 = -3.00, *P*Bonferroni = 0.0129; first- and second-order terms: *t*45 = -5.23, *P*Bonferroni < 0.0001; both compared to the intercept-only model; Figure 6A). No improvement in correlation was detected (first-order only: *t*45 = -0.99, *P*Bonferroni = 0.9777; first- and second-order: *t*45 = -0.49, *P*Bonferroni = 1; Figure 6A). Furthermore, no improvement in similarity was observed when including both the first- and second-order terms for the percentage of censored frames compared to only including the first-order term (RMSE: *t*45 = 0.02, *P*Bonferroni = 1; Pearson’s *r*: *t*45 = 1.75, *P*Bonferroni = 0.2589; Figure 6A). Finally, because pseudo-random frame censoring was shown to introduce a bias towards shorter INT values, we also compared the mean error across correction approaches. Both the intercept-only model and the model including a first-order term for the percentage of censored frames had mean error significantly different from zero (intercept only: *t*45 = -4.48, *P*Bonferroni = 0.0003; first-order term: *t*45 = 3.43, *P*Bonferroni = 0.0078; Figure 6A), indicating remaining bias in the corrected pseudo-randomly frame censored INT maps. No such bias remained in the corrected pseudo-randomly frame censored INT maps when including both the first- and second-order terms for the percentage of censored frames (*t*45 = -1.21, *P*Bonferroni = 1; Figure 6A). Furthermore, compared to the model including both the first- and second-order terms, the intercept-only model had greater negative bias (*t*45 = -11.30, *P*Bonferroni < 0.0001; Figure 6A) and the model including only the first-order term had greater positive bias (*t*45 = 46.86, *P*Bonferroni < 0.0001; Figure 6A). Together, these results suggest that including both the first- and second-order terms for the percentage of censored frames during group-level correction (i.e., using a quadratic polynomial for censored frames) eliminates a substantial portion of frame-censoring-induced underestimation bias.

**Figure 6.**
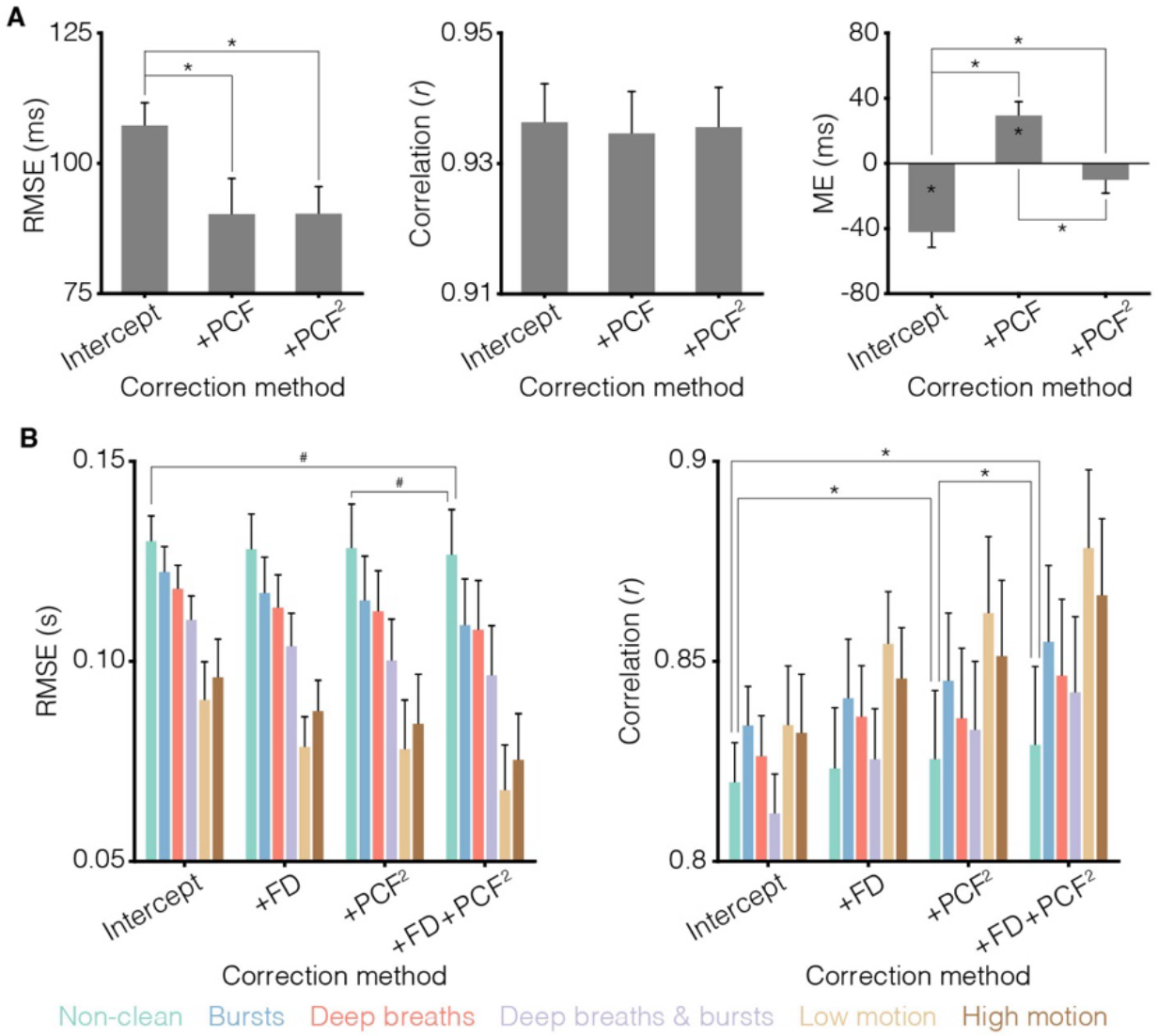
Group-level correction models can reduce the bias introduced by high-motion frame-censoring and residual effects of physiological and non-physiological artifacts. (A) Performance of group-level correction models on the pseudo-randomly frame censored runs.

We next sought to test this group-level correction approach in the non-clean data. Here, we also included the mean FD in non-censored frames in the group-level correction model.

Similar to the analysis using pseudo-random frame censored data, the group-level correction model including first- and second-order terms for the percentage of censored frames and the mean FD in non-censored frames improved similarity between the corrected non-clean INT maps and the clean INT maps (RMSE: *t*44 = -2.34, *P*Bonferroni = 0.0708; Pearson’s *r*: *t*44 = 3.94, *P*Bonferroni = 0.0009; both compared to intercept only model; Figure 6B). Meanwhile, no consistent improvements in similarity were observed for the group-level correction models only including mean FD (RMSE: *t*44 = -2.12, *P*Bonferroni = 0.1181; Pearson’s *r*: *t*44 = 2.95, *P*Bonferroni = 0.0150; both compared to intercept-only model; Figure 6B) or only including the first- and second-order terms (RMSE: *t*44 = -1.21, *P*Bonferroni = 0.6984; Pearson’s *r*: *t*44 = 2.89, *P*Bonferroni = 0.0178; both compared to intercept-only model; Figure 6B). Furthermore, the group-level correction model controlling for both the percentage of censored frames and the mean FD in non-high-motion frames outperformed the group-level correction model only including terms for the percentage of censored frames (RMSE: *t*44 = -2.38, *P*Bonferroni = 0.0431; Pearson’s *r*: *t*44 = 2.66, *P*Bonferroni = 0.0220), suggesting the need to control for residual motion in non-censored frames (Fig. 6B).

The inclusion of a linear term for the percentage of censored frames (PCF) and its square (PCF^2^) significantly improves the similarity (RMSE only; left) between pseudo-randomly frame censored and clean (non-censored) runs compared. Significant bias remains in the data corrected with either the intercept only model (negative bias) or the model including a linear term for PCF (positive bias) and the remaining bias is significantly less in the data corrected with the model including both linear and quadratic terms for the percentage of censored frames (right). Intercept: only includes an intercept; +PCF: includes an intercept and a linear term for the percentage of censored frames; +PCF^2^: includes an intercept, a linear term and quadratic term for the percentage of censored frames. (B) Performance of group-level correction models on the non-clean runs. The inclusion of terms for framewise displacement (FD), PCF, and PCF^2^ significantly improves the similarity between non-clean and clean runs. Intercept: only includes an intercept; FD: includes an intercept and a linear term for the mean FD in non-censored frames; PCF^2^: includes an intercept, a linear term and quadratic term for the percentage of censored frames; FD & PCF^2^: includes an intercept, a linear term for the mean FD in non-censored frames, a linear term and quadratic term for the percentage of censored frames.

## DISCUSSION

In this work, we sought to characterize the influences of respiration and head-motion artifacts on INT estimation, and the efficacy of high-motion frame censoring and GSR on mitigating these effects. Leveraging a subsample of the HCP-YA dataset carefully annotated for burst and deep breath respiratory events (Lynch et al., 2020), we first identified subjects with at least one rs-fMRI run with no respiratory artifacts and minimal head motion to serve as “ground-truth” clean runs. We then examined the impacts of bursts, deep breaths, and head motion on INT estimates using a within-subject approach that isolates artifacts from between-subject trait effects—e.g., individual traits that may relate simultaneously to increased head motion and neural alterations that can therefore confound the interpretation of fMRI data (Williams et al., 2022). We observed that non-clean, burst, burst and deep breath, and high motion runs showed general increases in INT, whereas deep breath and low motion runs showed small and non-significant differences from clean runs. We also observed that higher levels of head motion correlated with longer INT. Additionally, we used pseudo-random frame censoring to uncover INT estimation biases induced solely by frame censoring and established a method for correcting these effects in group-level regression analyses.

Importantly, the magnitude of artifactual influences on INT that we detected was comparable to reported effect sizes of INT relating to clinical status. Going forward, it will be crucial for clinical and cognitive studies of INT to carefully control for and contextualize results in terms of the potential influences of such artifacts.

Our observations show that well-established artifactual effects of head motion and respiration on other rs-fMRI measures extend to INT. Contextualizing these results to the greater rs-fMRI literature, previous work has demonstrated that rs-fMRI functional connectivity estimates are inflated for high-motion time points and high-motion subjects (Burgess et al., 2016; Power et al., 2014), as well as for respiration-related effects (Lynch et al., 2020). While functional connectivity measures index between-voxel similarity (as opposed to within-voxel similarity over time; i.e., temporal autocorrelation), the convergence between these results and our findings in INT suggest that head motion and respiration introduce shared (artifactual) signals that inflate both temporal autocorrelation and functional connectivity. To address these artifacts, we propose the use of high-motion frame censoring with Lomb-Scargle interpolation of censored timepoints, GSR, and a group-level correction for the amount of frames censored (and its square).

Frame censoring is a well-established tool for reducing artifact-related noise (Ciric et al., 2017; Parkes et al., 2018; Power et al., 2020; Power et al., 2014). However, recent computational work suggests that the estimation of INT is sensitive to data paucity, risking statistical biases that result in underestimation of timescales (Zeraati et al., 2022). Motivated by this work, we isolated the effects of frame censoring through a pseudo-random frame-censoring approach and found that frame censoring generally caused decreased INT throughout the brain, in line with Zeraati et al.’s findings. The convergence between Zeraati et al. and our empirical results suggests that our previous finding that higher motion (mean FD) was associated with shorter INT (Wengler et al., 2020) could indeed have been driven by partial frame censoring (rather than motion artifacts per se). Most robust to this effect was Lomb-Scargle interpolation of censored data. Previous work has shown that interpolation of high-motion frames reduces the amplitude of artifactual signal spread into adjacent timepoints during frequency filtering, but it has been recommended that interpolated frames should then be re-censored (Power et al., 2014). This re-censoring may be appropriate for functional connectivity analyses where correlations are performed between distinct voxels, but it disrupts the autocorrelation in the data, leading to decreased INT—apparent in our analyses where re-censoring was performed (“zeroing” in Fig. 4). Notably, we also showed that a regression-based correction aids in reducing frame censoring-induced biases in INT estimation. Importantly, correcting for the mean FD, the amount of censored frames, and the square of the amount of censored frames outperformed simpler models for INT bias correction. In practice, this correction approach can be implemented by controlling for these covariates in standard group-level statistical analyses.

While GSR is a somewhat controversial method in the rs-fMRI literature, it appears to be the most effective tool for eliminating the effects of respiration artifacts. Main concerns with GSR relate to the introduction of spurious anticorrelations in resting-state fMRI data (Anderson et al., 2011) and the potential removal of global neural signals of interest (Gotts et al., 2013). But GSR also offers unique advantages. Empirically, motion-based regression and models of respiration-related signals have been shown to be ineffective in fully removing global respiration-related signals (Power et al., 2017b), while GSR has been demonstrated to effectively eliminate the distinct effects of respiration (Lynch et al., 2020). In the present study, GSR significantly increased the similarity between non-clean and clean runs when used in conjunction with high-motion frame censoring, but not when used alone. Furthermore, GSR did not seem to introduce any artificial patterns in the fMRI INT maps (unlike for rs-fMRI connectivity measures), instead making them more similar to the “ground-truth” clean runs when combined with frame censoring (Figure 5). Therefore, our results indicate that GSR does not induce biases in INT estimation, and that it aids in reducing artifact noise when combined with high-motion frame censoring. We therefore recommend the inclusion of GSR in INT estimation pipelines.

Difficulties in comparing denoising strategies and optimizing artifact correction arise from the absence of a “noise-free” ground truth. Here, we leverage clean runs as an approximation of a within-subject ground truth. This approach avoids an important confound in between-subject approaches to probing artifacts: certain populations, e.g., neuropsychiatric populations, may move more on average and also have true neural differences with respect to other populations (Williams et al., 2022). The between-subject approach risks assuming all fMRI changes in high-motion individuals are artifactual and can therefore miss true neural differences. Aside from these designs, others have posited that simulations are the best approach to systematically evaluating the impact of artifacts since they circumvent the issue of establishing what is considered noise from empirical data (Uddin, 2017), although concerns raised for such approaches also highlight the low-dimensionality of simulated data utilized in previous work and their consequent limited resemblance to real fMRI data (Power et al., 2017a). The lack of a specific temporal structure for identifying neural signals of interest in rs-fMRI is an additional challenge for assessing denoising strategies (Power et al., 2020). Hence, an event-related approach (similar to task-fMRI) centered on common behaviors during resting-state scans, e.g., respiratory patterns and head motion (as done in the present work), can greatly aid in the selection of denoising strategies for resting-state data.

In conclusion, we recommend implementing GSR and high-motion frame censoring with Lomb-Scargle interpolation of censored data and performing group-level correction of the bias introduced by frame censoring for rs-fMRI INT studies. Furthermore, given the sensitivity of INT estimation to head motion and data quantity, and the inability to ensure complete elimination of potential biases, in addition we strongly encourage explicit post-hoc group-level analyses to definitively rule out the possibility that any INT effects (e.g., differences between a clinical and a non-clinical group) are simply due to differences in head motion, respiration, or amount of data, including but not limited to post-hoc analyses in matched subsamples. We hope that following these recommendations will contribute to a more rigorous and scientifically fruitful body of INT work around this important rs-fMRI measure.

## DATA AND CODE AVAILABILITY

All data analyzed in this study are part of the publicly available Human Connectome Project Young Adult dataset (Van Essen et al., 2013). Code used for this study are available from the corresponding authors upon reasonable request.

## AUTHOR CONTRIBUTIONS

**Andrew Goldberg:** Data Curation; Formal Analysis; Investigation; Methodology; Software; Visualization; Writing – Original Draft Preparation; Writing – Review & Editing. **Isabella Rosario:** Investigation; Project Administration; Writing – Original Draft Preparation; Writing – Review & Editing. **Jonathan Power:** Conceptualization; Methodology; Writing – Review & Editing. **Guillermo Horga:** Conceptualization; Funding Acquisition; Methodology; Resources; Supervision; Writing – Review & Editing. **Kenneth Wengler**: Conceptualization; Data Curation; Formal Analysis; Methodology; Software; Supervision; Validation; Visualization; Writing – Original Draft Preparation; Writing – Review & Editing

## ACKNOWLEDGEMENTS

Data were provided by the Human Connectome Project, WU-Minn Consortium (Principal Investigators: David Van Essen and Kamil Ugurbil; 1U54MH091657) funded by the 16 NIH Institutes and Centers that support the NIH Blueprint for Neuroscience Research; and by the McDonnell Center for Systems Neuroscience at Washington University.

